# Relations between hemispheric asymmetries of grey matter and auditory processing of spoken syllables in 281 healthy adults

**DOI:** 10.1101/2020.08.12.247841

**Authors:** Tulio Guadalupe, Xiang-Zhen Kong, Sophie E. A. Akkermans, Simon E. Fisher, Clyde Francks

## Abstract

Most people have a right-ear advantage for the perception of spoken syllables, consistent with left hemisphere dominance for speech processing. However, there is considerable variation, with some people showing left-ear advantage. The extent to which this variation is reflected in brain structure remains unclear. We tested for relations between hemispheric asymmetries of auditory processing and of grey matter in 281 adults, using dichotic listening and voxel-based morphometry. This was the largest study of this issue to date. Per-voxel asymmetry indexes were derived for each participant following registration of brain magnetic resonance images to a template that was symmetrized. The asymmetry index derived from dichotic listening was related to grey matter asymmetry in clusters of voxels corresponding to the amygdala and cerebellum lobule VI. There was also a smaller, non-significant cluster in the posterior superior temporal gyrus, a region of auditory cortex. These findings contribute to the mapping of asymmetrical structure-function links in the human brain, and suggest that subcortical structures should be investigated in relation to hemispheric dominance for speech processing, in addition to auditory cortex.

## Introduction

Roughly 85% of people have left-hemisphere language dominance, as assessed with functional magnetic resonance imaging (fMRI) (Mazoyer et al. 2014), Wada testing (Keller et al. 2018), or functional Transcranial Doppler Sonography (Knecht et al. 2000). Consistent with dominance of the left hemisphere for processing speech, most people also show right-ear advantage when identifying spoken syllables in the dichotic listening paradigm, i.e. when presented with different syllables into the two ears (Hugdahl 2011). The exact population proportion showing left-hemisphere dominance varies across these methods and according to the different threshold criteria applied, as well as experimental details such as the language-related task to be performed (Mazoyer et al. 2014). For the dichotic listening task specifically, the proportion showing right-ear advantage depends on factors including stimulus order, the number of trials, attentional aspects of the task, and the native language of participants (Westerhausen 2019; Bless et al. 2015b).

The structural basis of hemispheric language dominance remains unclear. At the population level, the human brain shows various left-right asymmetries of structure, including hemispheric differences of cortical surface area and thickness in inferior frontal and lateral temporal regions, which are important for language and/or auditory processing (Kong et al. 2018; Toga and Thompson 2003; Geschwind and Levitsky 1968). Asymmetries of regional anatomy around the sylvian fissure (i.e. the fissure that separates the frontal and parietal lobes from the temporal lobe) have a developmental origin in utero (Kasprian et al. 2011), and are therefore likely to arise from a genetically regulated program that favours hemispheric differences (Francks 2015; de Kovel et al. 2018; de Kovel et al. 2017; Ocklenburg et al. 2017).

However, perisylvian regions also show a high degree of variability between people in terms of sulcal and gyral anatomy and asymmetry (Tzourio-Mazoyer et al. 2018b). There is also considerable variation in functional laterality for language, with around 10-15% of people having a bilateral pattern without clear dominance, and roughly 1% having completely reversed, right-hemisphere language dominance (Mazoyer et al. 2014). In recent years it has become clear that the population variances of language-related structural and functional asymmetry are only weakly correlated (Tzourio-Mazoyer et al. 2018b; Zago et al. 2017; Josse et al. 2009; Keller et al. 2018; Ocklenburg et al. 2014; Tzourio-Mazoyer and Seghier 2016; Greve et al. 2013; Jansen et al. 2010), which raises the question of how tightly asymmetrical structure is related to asymmetrical function. For example, it is uncertain whether surface area asymmetries of superior temporal auditory cortex correlate with hemispheric language dominance (Greve et al. 2013), or only with variation in more restricted regional functional asymmetries during auditory language tasks (Tzourio-Mazoyer et al. 2018b). Regardless, these studies found only subtle structure-function associations, not involving substantially predictive relationships.

We have previously suggested (Kong et al.) that asymmetrical structure may indeed be coupled to function in the population-average form that characterises the majority of people, but when asymmetrical development of the embryonic brain is perturbed in a minority of people, different aspects of asymmetrical organization may become dissociated from each other as re-organization occurs. This would be consistent with weak correlations between measures of asymmetrical structure and function in the adult population. In this context, it is still informative to identify subtle associations between population variances in asymmetrical structure and function, as the anatomical regions implicated may be especially important for underlying functional asymmetry in most people. Random variation in the early embryo may be the most important source of individual differences in adult brain asymmetry, because despite the emergence of population-average brain asymmetries *in utero*, variation in brain and behavioural asymmetries tends to be only weakly heritable (Kong et al. 2018; Eyler et al. 2013; de Kovel et al. 2019b; Postema et al. 2020; de Kovel et al. 2019a; de Kovel and Francks 2019; McManus et al. 2013), and only slightly affected by early life factors such as birthweight, twinning or breastfeeding (de Kovel et al. 2019b; Postema et al. 2020; de Kovel et al. 2019a; de Kovel and Francks 2019; McManus et al. 2013).

As regards structural correlates of functional asymmetry measured by dichotic listening specifically, only a small number of studies have been performed. In one study of 29 healthy adults, macro- and microstructural properties of the arcuate and uncinate fasciculus, two nerve fibre tracts that link core language-related cortical regions, were linked to the extent of right-ear advantage in the dichotic listening task (Ocklenburg et al. 2014). Another study of 24 people with schizophrenia, and 25 controls, reported that better performance for the left-ear stimulus was associated with larger grey matter volume in the left frontal lobe, for the two groups combined (Nygard et al. 2013). A third study found that among 39 people with early onset schizophrenia, those with an absence of right-ear advantage had a reduction of left temporal lobe volume compared with those with normal right-ear advantage, and also compared to a set of matched controls (Collinson et al. 2009). While these findings are intriguing, the relatively low sample sizes warrant caution, as statistical power would have been low to detect subtle links between structure and function. Low statistical power in a study not only reduces the chance of detecting true effects, but also the likelihood that statistically significant results reflect true effects (Munafo and Flint 2010).

Here we performed the largest study to date aimed at finding relations between hemispheric asymmetries of auditory syllabic processing and of grey matter, based on MRI and dichotic listening data from 281 healthy adults. We took a brain-wide mapping approach, given that previous studies have not reliably limited the search space, and that hemispheric asymmetries of auditory processing may relate to broader network-level functional asymmetries (Ji et al. 2019). However, we also considered superior temporal regions of primary and secondary auditory cortex (Moerel et al. 2014) as a specific candidate set for focused analysis.

## Methods

### Participants

The BIG study was initiated in 2007 and comprises healthy volunteers, including many university students, who have participated in studies at the Donders Centre for Cognitive Neuroimaging, Nijmegen, The Netherlands (Franke et al. 2010). Participants underwent anatomical (T1-weighted) MRI scans as part of their involvement in diverse smaller-scale studies at the Donders Centre. All participants gave written, informed consent to participate in BIG. After inclusion of their scan in BIG, participants were re-contacted several times over the course of the subsequent years, to request that they complete online test and questionnaire batteries, one of which included the dichotic listening task (below). At the time of the current study, 643 participants had completed the relevant battery. After matching the available dichotic listening and neuroimaging data, and applying quality control criteria to both (specified below), in total 281 participants (180 females) were used in the dichotic-neuroimaging association analyses in this study. The mean age at MRI scanning for these 281 participants was 25.7 years old, SD 10.6 years, range from 18.1 to 69.0 years. The time that had passed between MRI scanning and dichotic listening ranged from 1.6 months to 15.3 years, mean 5.8 years (SD 3.7 years)(some scans were retrospectively included in BIG from as far back as 2003). The mean age was 31.4 years at the time of completing the online test battery, SD 11.1 years, range from 18.8 to 74.4 years.

Handedness was assessed with a questionnaire (a choice of strongly right-handed / moderately right-handed / ambidextrous / moderately left-handed / strongly left-handed, in Dutch, from which the ‘moderate’ data were then pooled together with the ‘strong’ data for each hand). Of the 281 participants with post-QC dichotic listening and MRI data, there were 10 left handers, 1 who indicated mixed hand preference, and 9 with missing handedness data. This rate of left-handedness (3.6%) is lower than the general population rate of roughly 10% in Northern Europe (de Kovel et al. 2019b; McManus 2009), as left-handedness was an exclusion criterion in some of the separate studies that contributed to the BIG dataset.

### Neuroimaging

Scanning was conducted using either a 1.5T Siemens Avanto or Sonata scanner, or a 3.0T Siemens TIM Trio, Skyra, Prisma, or Prisma-Fit scanner (see Table 1 for the numbers of participants by scanner). A variety of different scanning protocols was used (Supplementary Table 1). T1-weighted images were available for 293 participants who had post-quality-control dichotic listening laterality indexes (below). Twelve participants were excluded because of imaging quality: 6 during an initial quality control procedure to identify visibly large artefacts (e.g. caused by head motion) which would likely prevent brain structure from being properly delineated, 5 because of skull stripping failure, and 1 with a technical failure related to file reading. Brain structural data was analysed with FSL-VBM (Douaud et al. 2007), http://fsl.fmrib.ox.ac.uk/fsl/fslwiki/FSLVBM), an optimised VBM protocol (Good et al. 2001) carried out with FSL (v5.0) tools (Smith et al. 2004). First, structural images were brain-extracted and grey matter-segmented before being registered to the MNI 152 standard space using non-linear registration (Andersson et al. 2010). The resulting images were averaged and flipped along the x-axis to create a left-right symmetric, study-specific grey matter template. Second, participants’ grey matter images were non-linearly registered to this study-specific template and “modulated” to correct for local expansion or contraction due to the non-linear component of the spatial transformation. The modulated grey matter images were then smoothed with an isotropic Gaussian kernel with a sigma of 2.55 mm (full-width at half-maximum=6mm). Next, voxel-wise asymmetry index (AI) maps were calculated using the formula (L-R)/((L+R)/2) for each pair of leftright corresponding voxels, per participant.

**Table 1.** The numbers of participants per MRI scanner. See also Supplementary Table 1 for scanning protocols.

| *Scanner* | *Field Strength* | *Participants* |
| --- | --- | --- |
| Siemens Avanto | 1.5 T | 66 |
| Siemens Sonata | 1.5 T | 14 |
| Siemens TIM Trio | 3 T | 109 |
| Siemens Skyra | 3 T | 46 |
| Siemens Prisma | 3 T | 30 |
| Siemens Prisma-Fit | 3 T | 16 |
| **Total** | **-** | **281** |

### Dichotic listening

The dichotic listening task was performed at home via web-browser, implemented by the Delosis company (Delosis.com). Participants used either their own earphones, or earphones provided to them by mail. Participants were first presented with the test instructions and 4 questions dealing with the correct left-right placement of their headphones. For 3 of the questions, participants selected one of three possible options for the perceived location of a test tone (left ear, right ear or both). The 4th question was answered by moving a slider on a scale ranging from 0 (left) to 100 (right) for a tone presented equally to both ears. 410 participants correctly answered all four of the 4 setup questions (for the slider scale, responses between 30 and 70 were accepted as correct). Data from participants who had answered one or more of the four questions incorrectly were discarded. Participants were also presented with an explicit instruction that the “ga” syllable referred to the /g/ sound as used in common loanwords into the Dutch language, rather than the Dutch pronunciation (/x/ or /⍰/ which is not a stop-consonant).

The dichotic listening task involved 6 consonant-vowel syllables as stimuli. This format has been used previously in a functional imaging study (Hugdahl and Westerhausen 2016), and was also shown to be effective in a smartphone application outside a laboratory environment (Bless et al. 2015b; Bless et al. 2013). Nonetheless, our exact implementation involved new recorded stimuli that were suitable for Dutch first-language participants, and for integration into our software application.

The stimuli consisted of consonant-vowel (CV) syllables using the six stop-consonants: /ba/, /da/, /ga/, /ta/, /ka/ and /pa/. The syllables were spoken with constant intonation and intensity by a male native Dutch speaker, 32 years old. These syllables differ in their voicing and can be grouped in two categories. Voiced stop-consonant syllables have a short (S) voice of onset time (VOT), while voiceless stop-consonant syllables have a long (L) VOT. The voiced stop-consonant syllables (S), /ba/, /da/ and /ga/, had durations of 450, 467 and 506 ms respectively. The voiceless stop-consonant syllables (L), /ta/, /ka/ and /pa/, had durations of 305, 300 and 310 ms respectively. The syllables were normalized for loudness and each pair was temporally aligned for simultaneous release of the stop-consonants, using Audacity, v2.1 (https://audacityteam.org/). All syllable combinations were formed in both left-right and right-left orientations, resulting in a total of 36 pairs, i.e. 30 dichotic and 6 homonym pairs (Table 2).

**Table 2.**
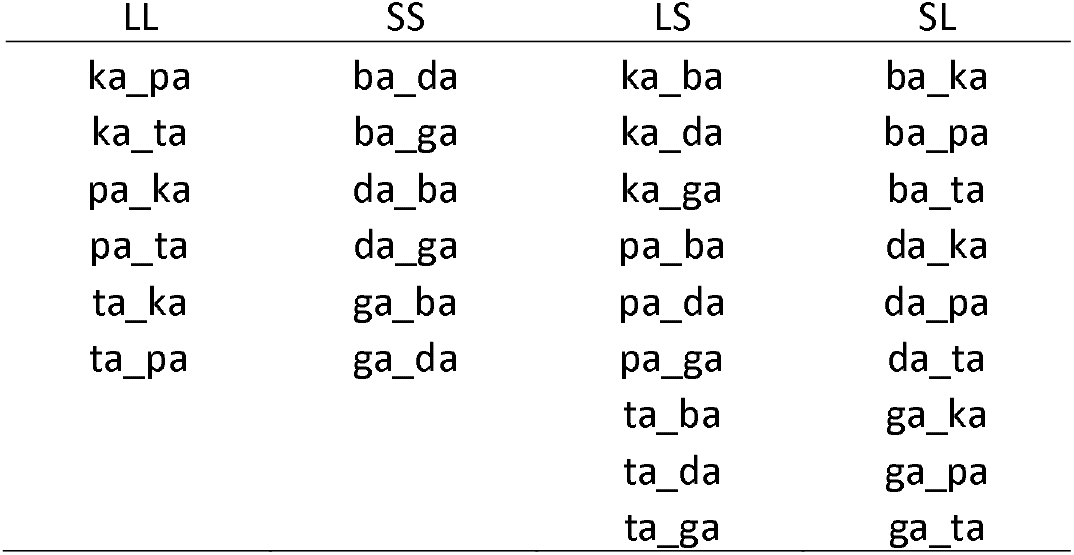
The 30 dichotic stimulus pairs, arranged according to voice onset time (VOT) categories. LL = long to both ears; SS = short to both ears; LS = long to left, short to right; SL = short to left, long to right.

| LL | SS | LS | SL |
| --- | --- | --- | --- |
| ka_pa | ba_da | ka_ba | ba_ka |
| ka_ta | ba_ga | ka_da | ba_pa |
| pa_ka | da_ba | ka_ga | ba_ta |
| pa_ta | da_ga | pa_ba | da_ka |
| ta_ka | ga_ba | pa_da | da_pa |
| ta_pa | ga_da | pa_ga | da_ta |
|  |  | ta_ba | ga_ka |
|  |  | ta_da | ga_pa |
|  |  | ta_ga | ga_ta |

Each of the syllable pairs was presented once. After each presentation, participants clicked on the button corresponding to the syllable they could best identify, from a choice of all six. A new pair was presented automatically after 4000ms. Correct responses were recorded when a syllable was correctly identified for either ear, and missing responses were considered incorrect. Participants were excluded if they had an error rate > 80% for dichotic trials, or an error rate > 50% for homonym trials, following a previously reported protocol (Bless et al. 2015b; Bless et al. 2013).

We computed average correct responses for the four different presentation categories: 1) when both ears were presented long-VOT syllables at once, 2) when both ears were presented short-VOT syllables, 3) when a long-VOT syllable was presented to the left and a short-VOT syllable to the right, and 4) when a short-VOT syllable was presented to the left and a long-VOT to the right (Table 2). We noticed that for syllable pairs of unequal voicing categories (i.e. LS and SL) correct responses were given overwhelmingly for the short VOT syllable, regardless of ear presentation (Figure 1; see the Discussion for an interpretation of this issue).

**Figure 1.**
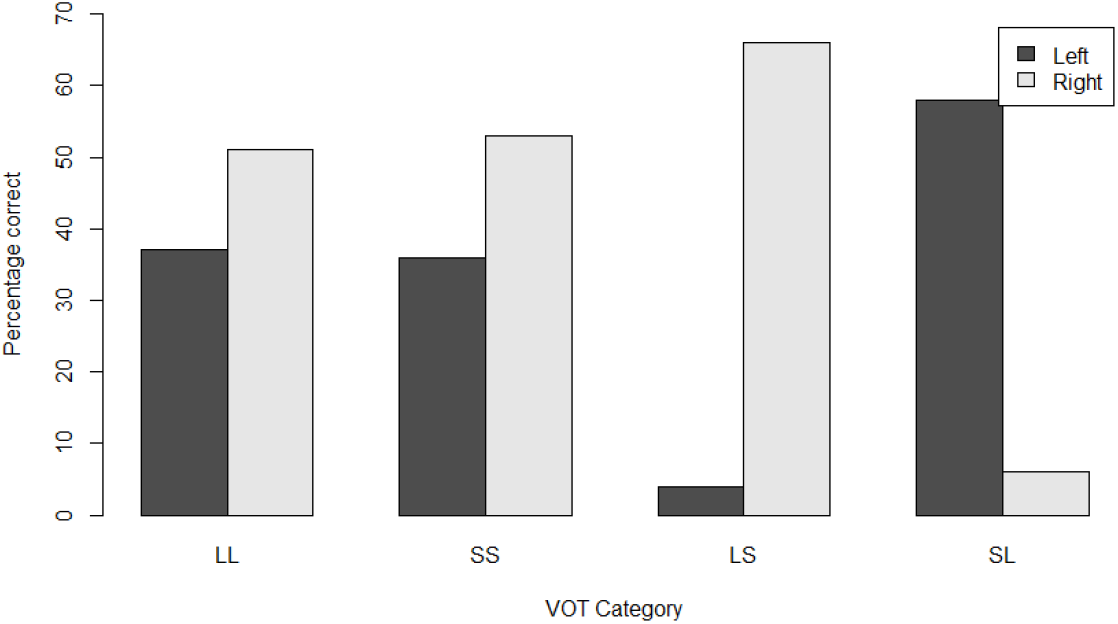
Average correct responses for the four different presentation categories by voice onset time (VOT) category. LL = long to both ears; SS = short to both ears; LS = long to left, short to right; SL = short to left, long to right.

We therefore computed the dichotic listening laterality index based only on presentations where both syllable pairs were of equal voicing group (i.e. LL and SS; Table 2). Laterality indices were calculated for each subject as the proportional difference in correct responses between right and left presented syllables: LI = (# correct Right – # correct Left) / (# correct Total), with positive scores denoting a right-ear advantage for identifying syllables.

### Voxelwise general linear model analysis

General linear modelling (GLM) was applied to examine the association between grey matter AI maps (voxel-wise AIs as dependent variables) and the dichotic listening laterality index (independent variable). Confound variables in the GLM were age at MRI scanning (continuous), years elapsed between scanning and dichotic listening (continuous), sex (binary), and scanner (categorical, as dummy coding, which also accounted for field strength differences; Table 1). We did not include handedness in the modelling, as only 10 participants were left-handed (see above) and there was no association of handedness with the dichotic listening laterality index (Left vs. Right: t = −0.93, p = 0.36; see also Discussion). Multiple comparison correction across the whole brain was performed on statistical maps using the *easythresh* program (voxel-level Z > 3.09, and cluster-level p < 0.05) implemented within FSL. Clusters were given post hoc anatomical labels according to the Harvard-Oxford atlases of cortical and subcortical brain regions (Goldstein et al. 1999; Frazier et al. 2005), and the cerebellar atlas from FSL (Diedrichsen et al. 2009).

In addition, we used a relatively loose threshold (voxel-level Z > 2.58 & cluster-level p < 0.05) to examine three neighbouring candidate regions of interest within our symmetric, study-specific grey matter template, using a voxel-wise probability threshold of 50% for belonging to a given region (according to left-hemisphere regional definitions within the Harvard-Oxford cortical atlas (Goldstein et al. 1999)). The regions were, from anterior to posterior along the superior temporal lobe: ‘Heschl’s gyrus’, ‘Superior Temporal Gyrus, posterior division’, and ‘Planum temporale’. The intention was to detect any potential trends of association that might indicate structure-function relations within auditory cortex, but that were too weak to detect using standard correction for voxel-wise multiple testing. However, Bonferroni correction was applied for multiple testing across the three regions (i.e., 0.05/3=0.017).

To further investigate each hemisphere’s contribution to the associations identified through the voxel-wise AI analysis (see Results), we performed post hoc analyses of the associations between the dichotic listening laterality index and the grey matter volumes at the homologous locations in the two hemispheres separately, for the peak coordinates per cluster, using Pearson correlation.

Finally, we repeated the voxel-wise AI association analysis after replacing the age at scanning as a covariate with the age at dichotic listening instead. The two ages were highly correlated with each other (r=0.94) so that minimal changes were expected in the regional associations with the dichotic listening laterality index.

## Results

The mean dichotic listening laterality index in the 281 participants with post-QC dichotic listening and MRI data was 19.6 (SD=35.1), which indicates a higher percentage of correct responses for syllables presented to the right ear, i.e. the expected right-ear advantage at the population level (Figure 2).

**Figure 2.**
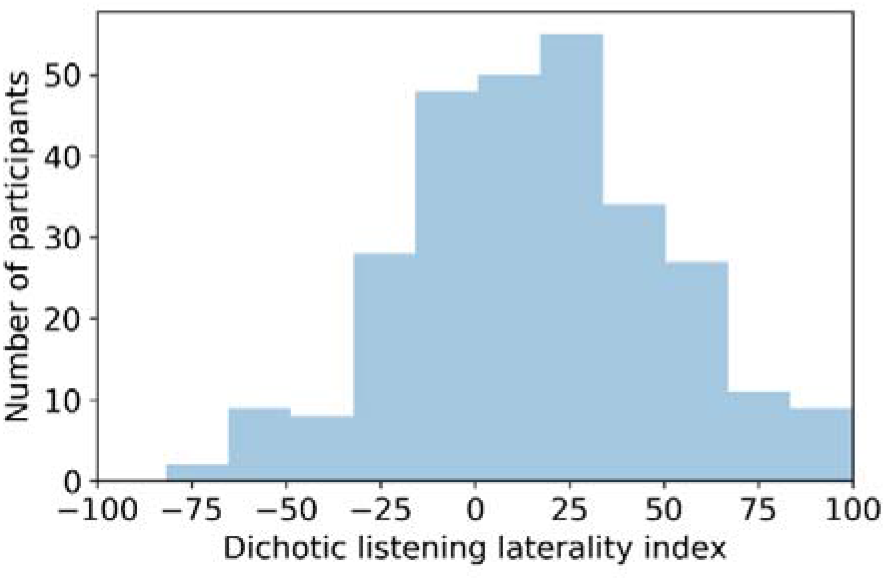
Frequency distribution of the dichotic listening laterality index in 281 participants with post-QC dichotic listening and neuroimaging data. Negative values indicate left-ear advantage, positive values indicate right-ear advantage.

In brain-wide analysis, the dichotic listening laterality index was significantly and positively associated with regional grey matter asymmetry in the amygdala (cluster size = 153 voxels, p = 0.0073; peak MNI coordinate: −14, −2, −20, Z = 3.98) and lobule VI of the cerebellum (cluster size = 134 voxels, p = 0.013; peak MNI coordinate: −26, −66, −18, Z = 3.85) (Figure 3A). In post hoc analysis of the corresponding unilateral grey matter volumes at the peak coordinates (i.e. in each hemisphere separately), both of these clusters showed greater right sided volumes in participants with increasingly atypical left-ear advantage, and no significant changes of the corresponding left-hemisphere volumes (Figure 3C).

**Figure 3.**
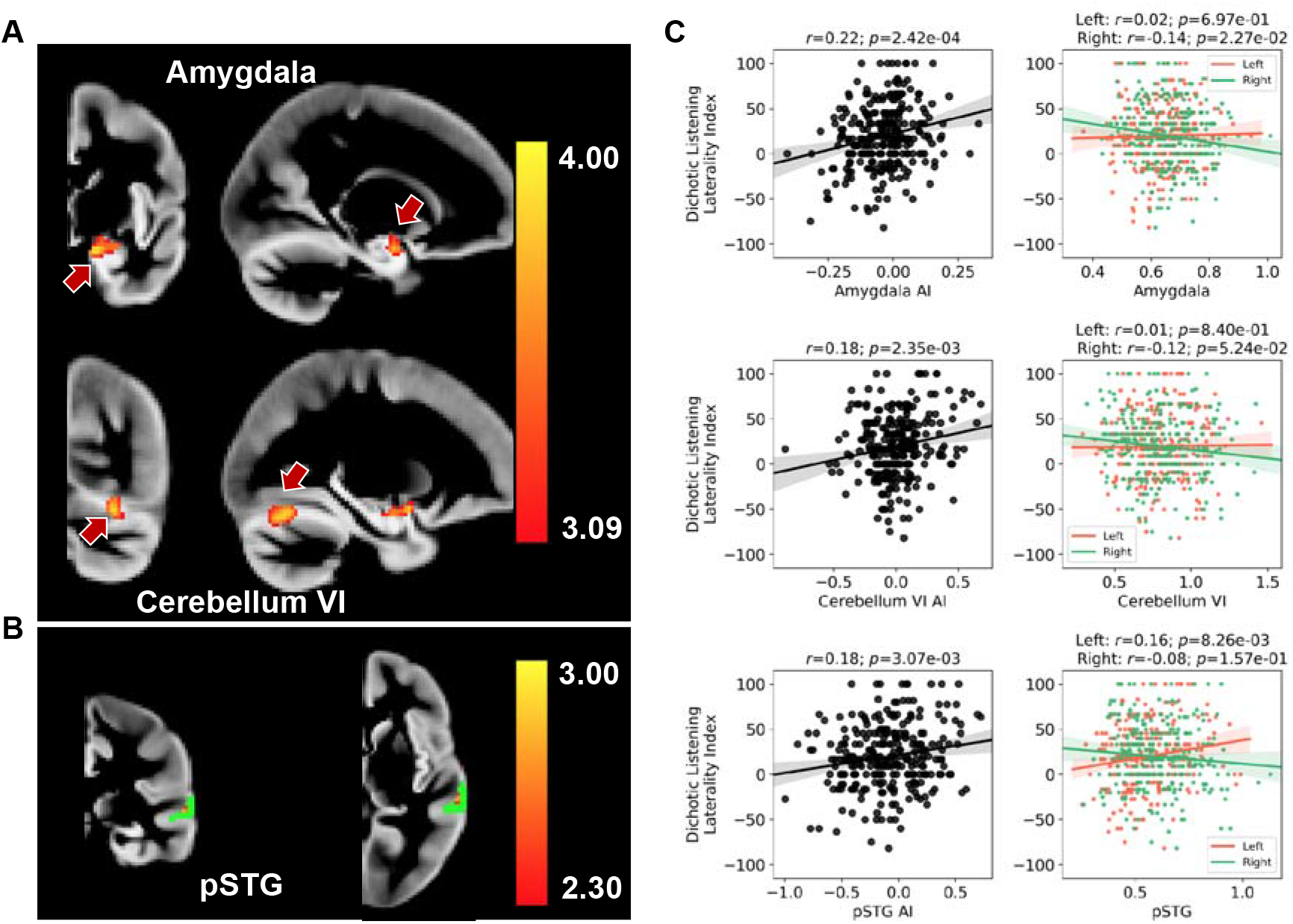
***A*** Two significant clusters associated with the dichotic listening laterality index, in voxel-wise grey matter asymmetry analysis; one in the amygdala, the other in cerebellum lobule VI. Both associations were in a positive direction (red-yellow indicates the association Z score). ***B*** Region-of-interest analysis of voxel-wise grey matter volume asymmetry in relation to the dichotic listening laterality index. The posterior superior temporal region of interest is shown in green (voxels with at least 50% probability of belonging to the region according to the Harvard-Oxford atlas), and a tentative cluster showing positive association is mapped in red-yellow (indicating the association Z score). ***C*** Directions of association between the dichotic listening laterality index and peak coordinate measures of grey matter volume, for the amygdala cluster (top), the cerebellum lobule VI cluster (middle), and the posterior superior temporal gyrus (pSTG) cluster (bottom). For each peak coordinate, results for voxel-wise grey matter asymmetry indexes (AI) as well as the corresponding unilateral left and right volume data are shown.

When focusing on three neighbouring regions of interest from the auditory cortex, using a relatively loose voxel-level Z threshold to detect any tentative trends of association (Methods), there was a small cluster on the posterior superior temporal gyrus (cluster size=3 voxels, corrected p = 0.014; peak MNI coordinate: −62, −30, 4, Z = 2.78; Figure 3B), within which grey matter asymmetry was positively correlated with the dichotic listening laterality index. In post hoc analysis of the corresponding unilateral grey matter volumes, smaller left-sided volume in this cluster was associated with increasingly atypical left-ear advantage, with no change on the right side (Figure 3C). There were no clusters identified within the Heschl’s gyrus or planum temporale regions of interest of the auditory cortex (p >0.10, uncorrected).

## Discussion

We performed the largest study to date of brain structural asymmetry in relation to auditory processing of spoken syllables. The dataset that we used was roughly one order of magnitude larger than previous studies that attempted to link dichotic listening laterality to variations in brain structure (see Introduction). In general, relations between structural and functional variability of the brain are subtle and complex (Chen and Omiya 2014; Tzourio-Mazoyer et al. 2018a; Batista-Garcia-Ramo and Fernandez-Verdecia 2018), and therefore studies of these relations are likely to benefit from larger samples than have often been used in the field.

In brain-wide voxel-wise asymmetry analysis, we found that two clusters, in the amygdala and cerebellum lobule VI, were significantly associated with the dichotic listening laterality index. These findings are striking insofar as they implicate regions located outside of the cerebral cortex, which is often considered as the primary seat of lateralized cognition in the human brain (Ji et al. 2019). As these are both original findings in our dataset, we remain cautious and suggest that further replication studies are needed. Nonetheless, there are other indications in the literature of the potential relevance of these structures to language-related cognition:

As regards the amygdala, data from stereo-electroencephalography (involving intracranial stimulation in patients with drug-resistant epilepsy) have shown functional connectivity with frontal and temporal regions important for language (Physiological Structural-Effective Connectivity Atlas: http://epi.fizica.unibuc.ro/atlas/). In addition, a recent fMRI study found the amygdala to be among regions showing left-lateralized activation during multi-modal, sentence-level task performance (Labache et al. 2019). Another recent fMRI study attempted to map relationships between subcortical structures and cortical networks, and found that the amygdala exhibits functional connectivity with the language network (Ji et al. 2019). This finding was flagged by the authors as unexpected and important to follow up in future research (Ji et al. 2019), as it does not currently have a clear or direct mechanistic interpretation. The amygdala is better known to be involved in emotion and motivation (Janak and Tye 2015).

Relatively more is known about the cerebellum in language-related processing (Desmond and Fiez 1998). The cerebellum has reciprocal connections with brain regions involved in phonological processing, i.e. the inferior frontal gyrus and lateral temporal cortex (Booth et al. 2007). We found a significant association between dichotic listening laterality and grey matter asymmetry specifically in cerebellum lobule VI, which was driven by a right-hemisphere increase of volume with atypical left-ear advantage. Language-related activation in the right cerebellum VI has been reported in language-task fMRI analysis (Stoodley and Schmahmann 2009), and is consistent with contralateral projections between the cerebellum and left-lateralized language network in the cerebral cortex (Jansen et al. 2005). Our data suggest that this reciprocal cerebellar-cortical functional laterality is linked to grey matter structural laterality in the cerebellum more noticeably than in the cerebral cortex, and point to a contralateral role of the cerebellum in syllabic auditory processing. Although tightly folded, the human cerebellar cortex was recently reported to have a surface area equal to roughly 4/5 of the neocortex, whereas in the macaque the cerebellar cortex is equal in surface area to only around 1/3 of the neocortex (Sereno et al. 2020). This suggests that the cerebellum has played an important role in the evolution of human cognition and behaviour, and language may be one adaptation to which the cerebellum contributes.

Developmentally, left-right differences of gene expression have been reported in the spinal cords and hindbrains of human embryos and fetuses, which may precede asymmetrical development of other brain regions (de Kovel et al. 2017). As a hindbrain structure, the cerebellum may therefore be a developmental source of laterality that affects other regions such as the cerebral cortex later on via asymmetrical, long-range connections. More research is required on hindbrain and subcortical roles in lateralized development and functioning of the human brain.

As regards auditory cortex, when using a relatively loose voxel-level Z threshold, we found a trend of association between the dichotic listening laterality index and grey matter asymmetry within the posterior superior temporal cortex, located next to Heschl’s gyrus and the planum temporale, where smaller left-sided volume was associated with increasingly atypical left-ear advantage. This direction of effect suggests that left-hemisphere dominance for auditory speech processing is favoured by greater left-hemisphere grey matter volume in a restricted region of auditory cortex. In a previous study contrasting 34 participants with left hemisphere language dominance and 21 with right hemisphere dominance (Greve, et al., 2013), analysis using surface-based registration across hemispheres and subjects found a small but significant cluster in the superior temporal gyrus, which overlapped with the planum temporale. The peak location reported in their paper (Greve, et al., 2013) does not match the cluster in our analysis, although the full extent of their cluster might have overlapped with ours (this is difficult to judge due to the different methods used). Our finding lends support to the existence of an anatomical correlate of functional language dominance within auditory cortex. However, functional laterality for auditory processing of spoken syllables cannot necessarily be taken to reflect broader hemispheric language dominance (Tzourio-Mazoyer et al. 2018b). Regardless, our finding is consistent with previous reports insofar as any superior temporal structure-function link is likely to be subtle, and not substantially predictive.

A previous fMRI study using dichotic listening in 104 participants found leftward asymmetries of activation in the posterior superior temporal gyrus, located adjacent to symmetrically activated areas (Westerhausen et al. 2014). Again, while the peak location of their asymmetrical cluster in the superior temporal gyrus did not co-locate with our anatomical cluster, it is possible that their broader cluster overlaps with ours, although it is not possible to be certain with the information given in the paper. We have deposited our full mapping results from the dichotic-asymmetry-VBM analysis in the Neurovault database (URL to be added on publication), in order to facilitate comparisons with future studies.

A limitation of our study was the time that elapsed between MRI scanning and dichotic listening, which ranged from 1.6 months to 15.3 years, and averaged 5.8 years. A necessary assumption of our study was therefore relative stability of brain structural and functional asymmetry over periods of this length in healthy adulthood, in a dataset of predominantly young adults. To partly address this potential issue, in addition to the age at scanning, we included the time difference between scanning and dichotic listening as a confound variable in our general linear modelling. Previous studies have indicated that age has significant but small effects on brain structural asymmetry (Fjell et al. 2015; Kong et al. 2018), such that age-related changes are likely to have limited impact during e.g. one decade of healthy adult life. Nonetheless, the statistical power to detect structure-function links in our study may have been reduced by this aspect.

Other limitations may relate to our Dutch-language, remote version of the dichotic listening task. Firstly, there was a substantial loss of data from 643 participants who attempted the task, to a final number of 293 with usable data (a further 12 were excluded due to quality control of MRI data). The loss of dichotic listening data occurred due to a combination of factors: failure to answer the setup questions correctly on left-right placement and balance of the headphones (the task was performed remotely at home), and high error rates which also incorporated missing responses. In principle, it is possible that loss of data was non-random, and that this led to biases in the dataset. However, there is no obvious reason to expect that brain asymmetrical structure-function links would be selectively sampled among those who provided usable data.

We also found that performance was strongly confounded with VOT (i.e. voice onset time) category, in the contrasts between short and long VOT stimuli, and we therefore excluded these pairs. This left twelve syllable pairs for calculating the dichotic listening laterality index, which may have contributed to error variance as the index was based on relatively few datapoints per participant. A previous study of the effects of VOT category in dichotic listening found the opposite pattern, i.e. long VOT stimuli were perceived dominantly, in contrasts with short VOT stimuli (Rimol et al. 2006). Our short VOT stimuli had longer overall durations than our long VOT stimuli (see Methods), and started before the long VOT stimuli, as all of the stimuli were aligned at the release of the stop consonant. These factors may have favoured dominant perception of short over long VOT stimuli in our study, regardless of the ear of presentation. To our knowledge, ours was the first Dutch implementation of consonant-vowel-syllable dichotic listening, and it has previously been found that performance on this task varies across languages (Bless et al. 2015a). More general limitations of dichotic listening for assessing hemispheric dominance for speech processing have been discussed extensively before (Hugdahl 2011; Westerhausen 2019).

As the MRI data in the BIG dataset came from multiple separate, smaller-scale MRI studies that were performed over a period of roughly fifteen years, there was substantial heterogeneity of scanners and scanning protocols (see Methods and Supplementary Table 1). We included ‘scanner’ as a confound variable in our general linear model analysis, which also adjusted for differences of field strength and other consistent properties of individual scanners. It is also likely that applying the denominator of the asymmetry index (L-R)/((L+R)2)) reduced the influence of scanning heterogeneity factors that impacted the measurement of both hemispheres equally. Nonetheless, if similar-sized studies could be carried out in the future using single scanners and scanning protocols, then reduced heterogeneity might improve the potential for detecting structure-function links. However, a doubt then arises about whether such findings would generalize across different scanners and protocols.

Rather than VBM, which measures grey matter volumes, future studies of brain structural relations to dichotic listening laterality may benefit from cortical surface-based approaches that separately measure surface area and thickness as distinct variables (Desikan et al. 2006). Improved mapping of asymmetry in this context may be achieved through hemispheric co-registration (Maingault et al. 2016; Greve et al. 2013). The precise mapping of VBM clusters is also an issue. For example, the cerebellar cluster identified in the present study overlapped at its upper edge with the nearby occipital fusiform gyrus, such that there may have been a degree of cortical contribution to this effect, but the resolution of the method was not sufficient to judge this with certainty (Figure 3). Other aspects of brain structure such as white matter tracts should also be investigated in larger datasets than used previously, through imaging modalities such as diffusion tensor imaging.

In our study, only 10 of the 281 participants with post-QC dichotic listening and brain MRI data were left-handed, due to left-handedness having been an exclusion criterion in some of the studies from which the BIG dataset was compiled (Guadalupe et al. 2014). There is an elevated rate of right-hemisphere language dominance among left-handers compared to right-handers (Mazoyer et al. 2014; Knecht et al. 2000; Van der Haegen and Brysbaert 2018), such that one of our ten left-handers would be expected to have right-hemisphere language dominance (Mazoyer et al. 2014). However, as the number of left-handers was too low for reliable statistical analysis, and we saw no association of the dichotic listening laterality index with handedness, we did not consider handedness further in our analysis. Handedness has also been found to be of limited relevance to cerebral cortical macrostructural brain asymmetry measures (Guadalupe et al. 2014; Kong et al. 2018). Larger studies, or studies which deliberately select for left-handed people, will be required to tease out potential relations between handedness, brain structure and auditory processing assessed by dichotic listening.

In summary, we have used MRI and dichotic listening data from 281 healthy subjects to contribute to mapping brain asymmetrical structure-function links, specifically with regard to grey matter and auditory processing of spoken syllables. The results identified novel associations involving the amygdala and cerebellum lobule VI, as well as a tentative cluster within auditory cortex. These findings motivate future studies of brain structural and language-related functional laterality that consider subcortical and hindbrain structures, in addition to the cerebral cortex.

## Acknowledgments

This research was funded by the Max Planck Society (Germany). This work made use of the BIG (Brain Imaging Genetics) database, first established in Nijmegen, The Netherlands, in 2007. This resource is now part of Cognomics (www.cognomics.nl), a joint initiative by researchers of the Donders Centre for Cognitive Neuroimaging, the Human Genetics and Cognitive Neuroscience departments of the Radboud University Medical Center (Radboudumc), and the Max Planck Institute for Psycholinguistics in Nijmegen.

## Data and code availability

Statistical maps before thresholding have been uploaded to NeuroVault (URL to be added on publication). Code for brain imaging analyses is available in GitHub (URL to be added on publication). Raw behavioural and imaging data were obtained from the BIG database and are available with the permission of the Cognomics Initiative (http://www.cognomics.nl/). Restrictions apply related to the privacy and consent of the research participants.

