## Supplementary information for "Relations between hemispheric asymmetries of grey matter and auditory processing of spoken syllables in 281 healthy adults"

**Supplementary Table 1: Scanning Protocols**

| **Protocol name** | **Numer_of Participants** | **Scanner** | **SliceThickness** | **PixelSpacing** | **AcquisitionMatrix** | **RepetitionTime** | **EchoTime** | **InversionTime** | **FlipAngle** |
| --- | --- | --- | --- | --- | --- | --- | --- | --- | --- |
| t1_mprage_sag_p2_iso_1.0 | 2 | TRIO | 1 | 1.0 1.0 | 0 256 256 0 | 2300 | 3.03 | 1100 | 8 |
| t1_mprage_sag_p2_iso_1.0 | 15 | PRISMAFIT | 1 | 1.0 1.0 | 0 256 256 0 | 2300 | 3.03 | 1100 | 8 |
| t1_mprage_sag_p2_iso_1.0 | 27 | PRISMA | 1 | 1.0 1.0 | 0 256 256 0 | 2300 | 3.03 | 1100 | 8 |
| t1_mprage_sag_p2_iso_1.0 | 40 | SKYRA | 1 | 1.0 1.0 | 0 256 256 0 | 2300 | 3.03 | 1100 | 8 |
| t1_mprage_sag_p2_iso_0.8 | 1 | SKYRA | 0.8 | 0.8 0.8 | 0 320 320 0 | 2300 | 3.15 | 1100 | 8 |
| t1_mprage_sag_iso_0.8 | 3 | SKYRA | 0.8 | 0.8 0.8 | 0 320 320 0 | 2300 | 3.15 | 1100 | 8 |
| t1_mprage_sag_iso_0.8LARGEFOV | 3 | PRISMA | 0.8 | 0.8 0.8 | 0 320 400 0 | 2200 | 2.64 | 1100 | 11 |
| golay_modified_erivdber2 | 1 | SONATA | 1 | 1.0 1.0 | 0 256 176 0 | 1660 | 2.48 | 750 | 8 |
| golay_modified | 4 | SONATA | 1 | 1.0 1.0 | 0 256 176 0 | 1660 | 2.02 | 750 | 8 |
| t1_mprage | 1 | TRIO | 1 | 1.0 1.0 | 0 256 256 0 | 2300 | 2.96 | 1100 | 8 |
| t1_mprage_TxRx_stad | 7 | TRIO | 1 | 1.0 1.0 | 0 256 256 0 | 2300 | 3.03 | 1100 | 8 |
| t1_mprage_sag_std | 13 | TRIO | 1 | 1.0 1.0 | 0 256 256 0 | 2300 | 3.03 | 1100 | 8 |
| T1_mprage_ADNI | 1 | TRIO | 1.2 | 1.0 1.0 | 0 256 256 0 | 2300 | 2.96 | 900 | 9 |
| t1_mpr_ns_optimal_contrast_PH | 1 | AVANTO | 1 | 1.0 1.0 | 0 256 256 0 | 2730 | 2.95 | 1000 | 7 |
| t1_mpr_ns_PH8_ADAM_GRAPPA2 | 4 | AVANTO | 1 | 1.0 1.0 | 0 256 256 0 | 2730 | 2.95 | 1000 | 7 |
| t1_mpr_ns_PH8_grappa2 | 7 | AVANTO | 1 | 1.0 1.0 | 0 256 256 0 | 2250 | 2.95 | 850 | 15 |
| t1_mpr_ns_PH8 | 19 | AVANTO | 1 | 1.0 1.0 | 0 256 256 0 | 2250 | 2.95 | 850 | 15 |
| t1_mprage_PH8_grappa2 | 30 | TRIO | 1 | 1.0 1.0 | 0 256 256 0 | 2300 | 3.03 | 1100 | 8 |
| t1_mprage_12Matrix_p2 | 1 | TRIO | 1 | 1.0 1.0 | 0 256 256 0 | 2300 | 3.03 | 1100 | 8 |
| t1_mprage_8_chan_grappa2 | 5 | TRIO | 1 | 1.0 1.0 | 0 256 256 0 | 2300 | 3.93 | 1100 | 8 |
| t1_mprage_sag_p2_iso_1.0_20ch_head | 2 | SKYRA | 1 | 1.0 1.0 | 0 256 256 0 | 2300 | 3.03 | 1100 | 8 |
| t1_mpr_ns_optimal_contrast | 1 | AVANTO | 1 | 1.0 1.0 | 0 256 256 0 | 2730 | 2.95 | 1000 | 7 |
| t1_mpr_ns | 8 | AVANTO | 1 | 1.0 1.0 | 0 256 256 0 | 2250 | 2.95 | 850 | 15 |
| t1_mpr_ns | 9 | SONATA | 1 | 1.0 1.0 | 0 256 256 0 | 2250 | 3.68 | 850 | 15 |
| t1_mprage_grappa2 | 10 | TRIO | 1 | 1.0 1.0 | 0 256 256 0 | 2300 | 3.03 | 1100 | 8 |
| t1_mpr_ns_optimal_contrast_32 | 1 | AVANTO | 1 | 1.0 1.0 | 0 256 256 0 | 2730 | 2.95 | 1000 | 7 |
| t1_mprage_32_grappa2_freesurfer_hubfon | 1 | TRIO | 1 | 1.0 1.0 | 0 256 256 0 | 2530 | 3.37 | 1100 | 7 |
| t1_mpr_ns_32 | 11 | AVANTO | 1 | 1.0 1.0 | 0 256 256 0 | 2250 | 2.95 | 850 | 15 |
| t1_mpr_ns_optimal_contrast_32ch | 13 | AVANTO | 1 | 1.0 1.0 | 0 256 256 0 | 2730 | 2.95 | 1000 | 7 |
| t1_mprage_32_grappa2 | 38 | TRIO | 1 | 1.0 1.0 | 0 256 256 0 | 2300 | 3.03 | 1100 | 8 |
| t1_structural | 1 | AVANTO | 1 | 1.0 1.0 | 0 256 256 0 | 2730 | 2.95 | 1000 | 7 |
| mp2rage_tr6_1mm_T1_Images | 1 | PRISMAFIT | 1 | 1.0 1.0 | 0 256 216 0 | 6000 | 2.34 | 0 | 0 |
